# Decoding the categorization of visual motion with magnetoencephalography

**DOI:** 10.1101/103044

**Authors:** Yousra Bekhti, Alexandre Gramfort, Nicolas Zilber, Virginie van Wassenhove

## Abstract

Brain decoding techniques are particularly efficient at deciphering weak and distributed neural patterns. Brain decoding has primarily been used in cognitive neurosciences to predict differences between pairs of stimuli (e.g. faces *vs.* houses), but how distinct brain/perceptual states can be decoded following the presentation of continuous sensory stimuli is unclear. Here, we developed a novel approach to decode brain activity recorded with magnetoencephalography while participants discriminated the coherence of two intermingled clouds of dots. Seven levels of visual motion coherence were tested and participants reported the colour of the most coherent cloud. The decoding approach was formulated as a ranked-classification problem, in which the model was evaluated by its capacity to predict the order of a pair of trials, each tested with two distinct visual motion coherence levels. Two brain states were decoded as a function of the degree of visual motion coherence. Importantly, perceptual motion coherence thresholds were found to match the decoder boundaries in a fully data-driven way. The algorithm revealed the earliest categorization in hMT+, followed by V1/V2, IPS, and vlPFC.

## Introduction

In natural environments, coherent motion is a vital sensory cue that helps the brain individuate objects in the world. Seminal neurophysiological work has described neurons in the middle temporal (MT) lobe of monkeys that were selective to the coherence^1^ and the direction^2^ of visual motion. During a perceptual classification task, direction-selectivity can be decoded from the activity of neural populations in MT^3, 4^. As visual motion processing relies on neural population codes, it is amenable to non-invasive functional human brain imaging such as fMRI or magnetoencephalography (MEG). Supervised learning techniques such as Multivariate Pattern Analysis (MVPA) are increasingly successful at characterizing where and when the neural analysis of stimuli such as visual orientation, motion direction or object classification is being realized^5–13^.

In one of the earliest fMRI studies using MVPA, the direction of motion was successfully decoded from hMT+ (human analog of MT) activity^14^) but also, and surprisingly, from visual cortices V1, V2, V3 and V4^15^. The successful decoding of visual motion in V1, V3 and hMT+ has since been reported several times^5, 15–18^. Visual motion decoding in lower visual areas has been functionally interpreted as an indication of feature-based attention when required by the task^15^ and as an effect of top-down modulation of early visual areas for conscious perception^16^. However, whether brain decoding using MVPA captures the selectivity of neural populations has been a subject of debate on the interpretational weigh given to decoding^11, 19–21^. Relevant to the current study, recent fMRI work has suggested that the sources of decoding in early visual areas may reflect the perceptual priors and biases of motion direction computation^22^.

To disambiguate the functional role of different brain regions in motion selectivity, characterizing the temporal unfolding of pattern classification within and across visual regions could be helpful. Here, we thus exploited the temporal sensitivity of MEG to find the latency at which sufficient information had been integrated to reach a stable classification boundary^23–25^. 36 participants were recorded with MEG while performing a visual motion coherence discrimination task in which two intermingled populations of visual dots (red and green random-dot-kinematograms) moved randomly on the screen until one of them moved more coherently than the other^26^ (Fig. 1-A). Participants were asked to report which of the two populations became most coherent over time. Seven motion coherence levels were tested and a novel multivariate decoding approach combining ridge regression and a ranking metric was developed. Contrary to classical decoding approaches based on binary classifiers such as support vector machines (SVM), a single decoder was estimated for all coherence levels, allowing robust parameter estimation despite high dimensional data. The ranking metric allowed taking into account the fact that visual motion coherence was an ordered variable^27, 28^. This novel decoder was applied to brain activity recorded at the sensor level and to cortically-constrained source estimates. Using this decoding technique, we report the categorization of two separate brain states as a function of the degree of visual motion coherence. The categorization boundary matched participants’ behavioral outcomes. Our results suggest that incorporating such decoding methods may be suitable to address questions relevant to predictive coding and perceptual decision-making.

**Figure 1.**
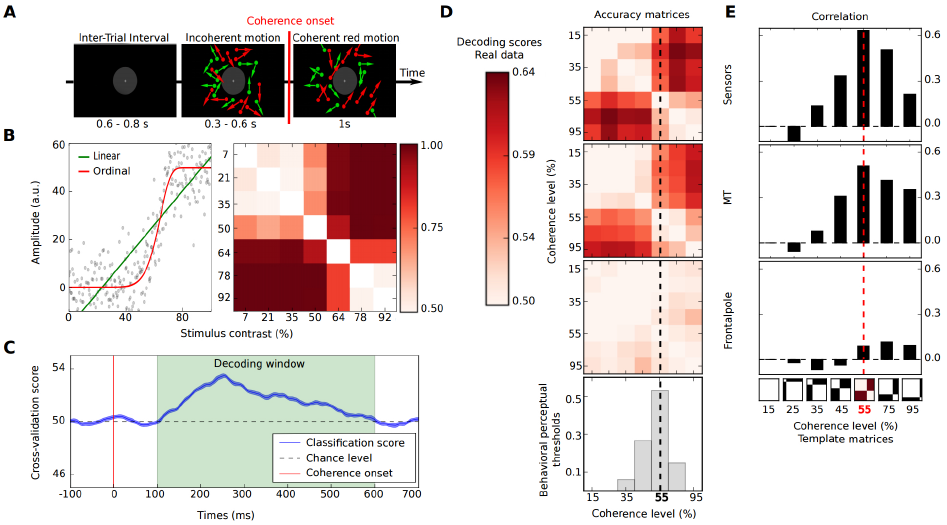
Categorization Decoding. a) One experimental trial in which participants discriminated which of the red or green cloud of moving dots was most coherent. b) Left: simulated data (gray) were best modeled by ordinal (red) than by a linear (red) fit. Right: similarity matrix providing a score of the decoding performance for each pairwise comparisons. c) Significant time-resolved decoding of visual motion coherence levels were found 100 to 600 ms (green) post-stimulus onset. d) Grand-average (n=36) similarity matrices in sensors (top), hMT+ (middle) and frontal-pole (bottom) for the selected time window. Distribution of behavioral perceptual thresholds (gray histogram) and the mean (dashed line). e) Correlation scores between each template and similarity matrix (black histograms) and likeliest boundary decoded from MEG data (dashed red line).

## Methods

### Participants

Thirty-six participants took part in the study (16 females, mean 22.1 +/- 2.2 y.o.). All were right-handed, had normal hearing and normal or corrected-to-normal vision. Prior to the experiment, all participants gave a written informed consent. All methods were carried out in accordance with relevant guidelines and regulations and by a named NeuroSpin (Gif-sur-Yvette, France). The study was conducted in agreement with the Declaration of Helsinki (2008) and was approved by the Ethics Committee on Human Research at Neurospin (Gif-sur-Yvette, France).

### Experimental design

The MEG session consisted of twelve experimental blocks alternating between rest and task ^26^. Here, we solely focused on the main experimental task blocks in which participants’ performance on a visual motion coherence task was being assessed. During the task, one trial started with the presentation of a fixation cross followed by two intermixed clouds of dots or Random Dot Kinematograms (RDKs) (red and green) whose motion was fully incoherent. After a variable interval of 0.3 to 0.6 s, one of the two RDKs became more coherent than the other (Fig. 1-A). Participant had to determine by button press which of the red or green RDKs became more coherent. Seven possible levels of visual motion coherence were tested (15%, 25%, 35%, 45%, 55%, 75%, or 95%), randomly assigned to a colour and to a direction. Each participant was tested with 28 trials per visual coherence level.

### Visual stimuli

The red and green RDKs were individually calibrated to isoluminance. To prevent local tracking of dots, a white fixation cross was located at the center of a 4° gray disk mask. RDKs were presented within an annulus of 4°-15° of visual angle. Dots had a radius of 0.2°. The flow of RDKs was 16.7 dots per deg^2^ × sec with a speed of 10°/s. During the first 0.3 to 0.6 s of a given trial, both RDKs were incoherent (0% of coherent motion). The duration of the incoherent phase was pseudo-randomized across each trial in order to increase the difficulty of the task by preventing participants’ expectation of the temporal onset coherent motion. After the incoherent phase, one RDK became more coherent than the other for one second. The direction of coherent dots was comprised within an angle of 45°-90° around the azimuth. 50% of the trials were upward coherent motion and the remaining 50% of the trials were downward coherent motion. At each frame, 5% of all dots were randomly reassigned to new positions and incoherent dots to a new direction of motion. Dots going into collision in the next frame were also reassigned a new direction of motion.

### Psychophysical analysis

The performance of each individual was averaged as a function of the seven degree of visual motion coherence of the stimuli, irrespective of colour or direction of motion. The coherence discrimination threshold was set to 75% of correctness for each individual’s data, as typically used in a two-alternative forced choice (2-AFC) paradigm, forcing participants to adopt the same decision criterion for all stimuli^29^. Here, the 75% detection threshold corresponds to chance level. They were then separately fitted to psychometric functions with the maximum-likelihood methodology (Psignifit^30^) which provided valid estimates of perceptual thresholds on a per individual basis (more details in ^26^).

### MEG pre-processing and source reconstruction

All data pre-processing and source-imaging were done according to well accepted MEG guidelines^31^. Signal-Space-Separation (BS) was performed on raw data using Maxfilter (Elekta-Neuromag^32^) to compensate for external magnetic interferences. MEG data were band-pass filtered (2 to 45 Hz), down-sampled to 250 Hz and epoched from -100 ms to 1000 ms relative to the onset of RDK coherence. Trials that were contaminated by artifacts were rejected (e.g. peak-to-peak amplitude difference above 150 microvolts in EOG data) leaving 89% of trials considered to have an appropriate signal-to-noise ratio. The cortically constrained source reconstruction was done using the dSPM method following the guidelines of the MNE software^33^. The entire pre-processing was done using MNE^34^.

### MEG decoding

Decoding generally consists in predicting a target variable *y* from one pattern of brain activity 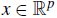
 among all possible patterns or brain states. When the target can take a finite number *K* of possible values, like a multi-class classification problem, one has that 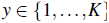. Here, when *x* were MEG signals, *p* was the number of channels and time points used for the prediction. When *x* was the amplitude of cortical sources, *p* corresponded to the number of source locations. The first goal of this study was to estimate how well each pair of visual motion coherence level could be discriminated against each other. Considering that multi-class classification approaches do not take into account the ordinal nature of the target to predict, indeed predicting 1 instead of 7 is as bad as predicting 1 instead of 2 although the mistake is obviously smaller in the second case, we instead built a decoder which could yield high pattern classification accuracy for distinguishable coherence levels, and low pattern classification accuracy for nearby levels of visual motion coherence which were perceptually hard to differentiate (cf. next two sections on method).

The second goal of the study was to find whether separate categorical brain states (two or more) emerged following the presentation of the stimuli as a function of the seven levels of visual motion coherence. Specifically, the task of participants consisted in deciding whether the red or the green cloud of dots was most coherent as a function of coherence level. One working hypothesis was thus that at least one boundary delimiting a possible threshold between the neural activations induced by low vs. high coherent motion would be found during decoding.

To address this question, we opted out of a regression model estimated jointly for all levels of coherence, and combined it with a ranking metric adapted to discrete and ordered targets. Although an alternative approach could have consisted in testing the incoherent portion of the stimuli against each level of visual coherence, this would have lead to a strongly imbalanced training dataset (i.e. 196 incoherence trials for 28 trials per level of coherence) which is heavily problematic for MVPA classification approaches^35^. Specifically, with this formulation of the decoding, an inaccurate model which always predicts incoherence instead of coherence would have 85% of accuracy due to the imbalanced dataset. The ranking technique proposed here does not suffer from such class imbalance considering that a single regression model was learnt for all coherence levels, and the ranking metric employed yielded 50% accuracy levels in spite of the low number of trials.

We now describe in detail the regression model employed.

### Regression model

Due to the limited number of data points available for learning, and the high dimensional nature of the neuroimaging data, we used a linear model following the standard approach in MVPA studies^5, 11, 23^. The target values 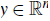 here provided for the *n* data points available for statistical inference, were derived from a linear combination of data 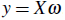 where 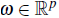 was a weight vector and *X* was a *n*-by-*p* data matrix. The value *n* here corresponded to the number of stimuli presentations, a.k.a. single trials or epochs. For each i*^th^* observation, the target 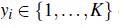 could take *K* different values: in this study, *K* = 7 corresponded to the seven levels of visual motion coherence defining the number of classes. Again, a multi-class classification approach could have been adopted, yet this strategy would have ignored that target values were ordered. For instance, decoding the 5th instead of the 2nd level of motion coherence is worse than predicting the 3rd level of motion coherence instead of the 2nd one. This is an information that a multi-class linear SVM model could not exploit. An SVM would also estimate *p × K* parameters instead of *p* which would naturally increase the risk of overfitting and reduced the interpretability of the results. Instead, we chose a ridge regression method, and evaluated the predictive performance with a metric tailored for ordinal problems. The ridge regression model was defined as the solution to the convex optimization problem:

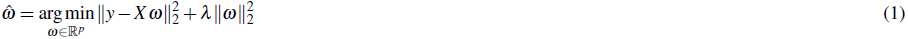

The ridge regression model is a popular approach whose practical success is due to fast estimation, robustness to noise and limited sensitivity to rough tuning of the parameter *λ*. Indeed results obtained by ridge regression are known to be far less sensitive to the choice of *λ* parameter compared to sparse estimators such as Lasso. In our experiments, *λ* was the same for all subjects^36^.

Decoding was performed on a per individual basis using all epochs. The 204 gradiometers and different time windows were tested: for example, for the time window ranging from 100 to 600 ms, the dimensions of the data were the number of samples *n* = 196 (at most, 28 trials × 7 coherence levels) depending on the number of dropped epochs times the number of features *p* = 204 × 126 ~ 2.5 × 10^4^ where the temporal window ranging from 100 ms to 600 ms contained up to 126 samples. The performance of the method was evaluated with a 10-fold stratified cross-validation which preserved the percentage of samples for each class or motion coherence level in each fold.

Decoding was also performed on source-reconstructed data in bilateral regions of interest (ROI) previously reported as being implicated in the task^26^. In source-space, the dimensions of the data were *n* = 196 at most and, for instance, *p* = 126x117~10^6^ depending on the size of the ROI (here, 117 dipoles in the ROI).

Following estimation of the ridge regression model, a ranking metric was then employed to quantify the model performance while taking into account that the targets have a natural order.

### Assessing decoding performance with pairwise ranking metric

Although ridge regression preserves the order of the target variables, it does not provide a relevant metric for the evaluation of the success rate of the decoder with an ordered set of categories. When using a linear regression model, the mean square error (MSE) is the natural performance metric. Yet, in high dimensional settings with a limited number of samples (*n* ≪ *p*) as we are dealing with here, MSE is a poor metric. In order to reduce the variance of the estimated coefficients, high values of *λ* were used causing a strong amplitude bias on the coefficients and a poor performance when measured using MSE. Performance evaluated with MSE was also affected in the presence of a bimodal state as illustrated in Fig. 1-B. Note that this strong bias problem is what motivates certain authors to use a Pearson correlation as measure of performance rather than the MSE, although MSE is natural when using ridge regression^37^.

To leverage the ordinal nature of the target values *y*, we quantified the performance in terms of ranking, where we tested the ability of the decoder to properly order pairs of samples, trials, based on the target to predict^27, 28^. The ranking metric consisted in comparing the real values of *y* and the predicted ones. Let us consider two trials from the validation dataset with 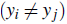 and where (*y_i_; y _j_*) denote their associated labels.

Let 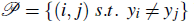 be the set of pairs with different labels. One quantifies the prediction accuracy *Acc* with the percentage of correct orderings for pairs of trials:

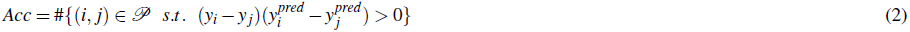

For each pair of trials, there were two possible options and the chance level was therefore 50%. This quantity is related to Kendall’s rank correlation metric^38^ which can be seen as a non-parametric correlation measure. To go beyond average accuracy, a key insight of this work was to inspect for which pair of trials the decoder made a mistake. For this, we thus defined a 7-by-7 similarity matrix *M*:

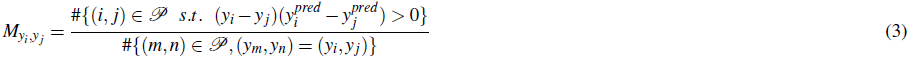

Each *M_i;j_* was a value between 0 and 1 that told us how well we could distinguish the level *i* from the level *j*, 1 being the best; inversely, if the level *i* was similar or close to the level *j*, this decoding value would be close to chance level 0.5. The matrix was symmetric since comparing the levels *i* and *j* or *j* and *i* provides the same score. Such matrices, that can be seen as confusion matrices adapted for our pairwise ranking metric, are presented in Fig. 1-D.

### Criteria for decoding categorization

Template matrices were defined for the discrete values of theoretically possible categorization into two brain states driven by the motion coherence levels, namely: 15%, 25%, 35%, 45%, 55%, 75%, or 95%. Each matrix had an on/off pattern at a given threshold (e.g. 55%) with values of 0.5 (off) or 0.65 (on). An example is provided in the black matrices of Fig. 1-E. The correlation between the empirical matrices (fully based on MEG data) and all the possible template matrices as defined above, thus provided the selection criterion to decode a categorization pattern at a specific threshold. Specifically, for each empirical similarity matrix, the template matrix yielding the highest correlation score was considered a good predictor of the participants’ motion coherence thresholds eliciting the choice boundary from MEG data indicated as a dashed vertical line in Fig. 1-D.

## Results

### Modeling of simulated data as proof of concept

First, we modeled typical behavioral profiles observed during a perceptual discrimination task by using simulated data (e.g. ranging from 7% to 92% of coherence). The modeling allowed validating the use of an ordinal model which fitted better the data than a linear model (Fig. 1-B, left panel). As detailed above, the simulated trials were decoded using cross-validation by fitting a ridge regression to the training data and evaluating the performance of the model on all possible pairwise combination of test trials. The similarity matrix (Fig. 1-B, right panel), which represents the predictive power in distinguishing two coherence levels, was evaluated with a 10-fold stratified cross-validation method. Each entry in the similarity matrix shows how similar each coherence level is to another one; alternatively, each entry can also be interpreted as how well one coherence level can be distinguished from another using a linear multivariate statistical model. All pairwise comparisons given in the similarity matrix built an anti-diagonal pattern: the lighter blocks in the similarity matrix were coherence levels for which no differences in brain responses could be captured yielding a decoding score at chance level; conversely, the darker blocks (red) captured high decoding accuracy scores for which brain responses highly differed between two coherent motion *e.g.*, brain responses to 7% coherent motion were highly distinguishable from those obtained during the presentation of 92% coherent motion. When comparing the neighboring levels 64% and 78% in (Fig. 1-B, right panel), the high accuracy of decoding demonstrated a difference in brain activity patterns, reflecting a discontinuity in the activation profiles despite a progressive change in the visual motion coherence levels. The observed discontinuity or edge located between 50% and 64% of visual motion coherence revealed the presence of a categorical boundary.

### Spatial selectivity of decoding categorization

The appropriate time window for best decoding performance was established using time-resolved cross-validation techniques^24^. The overall best decoding performance was obtained for latencies ranging from 100 ms to 600 ms post-motion coherence onset as illustrated in (Fig. 1-C). The decoder was applied to MEG data in this time window on a per individual basis. Similarity matrices scored how well pairs of visual motion coherence could be distinguished, and then ordered, on the basis of brain activity. Fig. 1-D reports the similarity matrices computed on grand-average MEG data (n = 36 participants). Similarity matrices obtained for the MEG sensors (gradiometers) are reported in the top panel. Similarity matrices obtained for source-reconstructed estimates in the ROI hMT+ and in a control region “frontal pole” are provided in the middle and bottom panels, respectively.

The similarity matrices obtained in sensor and hMT+ data showed two distinct categories as an anti-block-diagonal patterns: two light blocks of decoding score at chance level (~50%) for close coherence levels (low levels: 15%-45% against themselves, high levels: 55%-95% against themselves), and two dark blocks of decoding score nearing ~65% for coherence levels that were apart, namely 15%-45% against 55%-95%. These results conform with the notion of perceptual categories, namely: visual motion coherence levels 45% and 55% were close from the point of view of the coherence level in visual stimulation, but distant in perceptual space with the former most likely classified incoherent and the latter as coherent. The two brain states thus defined by the similarity matrix are compatible with categorical classification of the stimuli in this task. Specifically, visual motion coherence stimuli could either elicit a pattern consistent with not detecting the coherent signal in the display and not discriminating within the ensemble of stimuli whose coherence could not be detected (below the boundary) and detecting the coherent signal in the display but not discriminating within the ensemble of stimuli whose coherence could be detected (above the boundary).

To further investigate the link between brain activity at the single trial level and behavioral outcomes, we systematically compared the boundary delimited by the decoding approach with the perceptual threshold obtained from psychometric fits. The mean perceptual threshold was obtained in the task from the previous study^26^ and shown here in the histogram over the 36 subjects (Fig. 1-D, bottom panel). The emerging categorical boundary at 45-55% of visual motion coherence in both sensors and hMT+ (but not frontal pole) matched well the mean perceptual threshold observed behaviorally (black dotted line; Fig. 1-D).

To establish a quantitative criterion for this observation, template matrices were constructed to model each theoretically possible perceptual threshold. Each template matrix was then correlated with each of the decoding similarity matrices obtained from empirical measurements (Fig. 1-E). The aim was to find the peak of the correlation between the template threshold and the emerging boundary. This procedure, which is similar in spirit to the Representational Similarity Analysis (RSA) approach^9,39^, insured that the decoding similarity matrix was not forced to look like any specific template matrix. The quantitative metric confirmed our qualitative assessment (Fig. 1-D). Specifically, the peaks of the correlations were found for template matrices corresponding to a mean perceptual threshold of 55% in both MEG sensors and in source-reconstructed hMT+; the control ROI showed no selectivity.

### Temporal accumulation selectivity of categorization decoding

The spatiotemporal sensitivity of source-reconstructed MEG data was exploited to test at which latency sufficient information had been integrated to reach a reliable and stable classification pattern. To explicit the choice of the cumulative time window range, Fig. 2-A shows the grand average time course in response to the seven motion coherence levels over the 36 subjects in hMT+. As can be seen (Fig. 2-A) and as previously reported^26^, main differences were located at these latencies although no clear categorization were visible in the time response. For this, scoring was established in a temporally cumulative manner from 100 ms post-motion coherence onset on by adding the consecutive 50 ms time window to each previous one (Fig. 2-B) until 450 ms. The decoder was applied to sensors and to source estimates in the regions of interest as well as additional cortical sources known to be involved in the task^26^, namely: hMT+ and the control region frontal pole but also the medial primary and secondary visual cortices (V1/V2), the intraparietal sulcus (IPS) and ventrolateral prefrontal region (VLPFC) (Fig. 2, bottom left). In Fig. 2-B, in which all similarity matrices are reported, two brain categories of coherence levels seemed to emerge. As one of the focuses was to link the decoding to the behavioral data, the black dotted lines illustrated the known average perceptual threshold to find how well it fitted with the boundary found in the similarity matrices.

**Figure 2.**
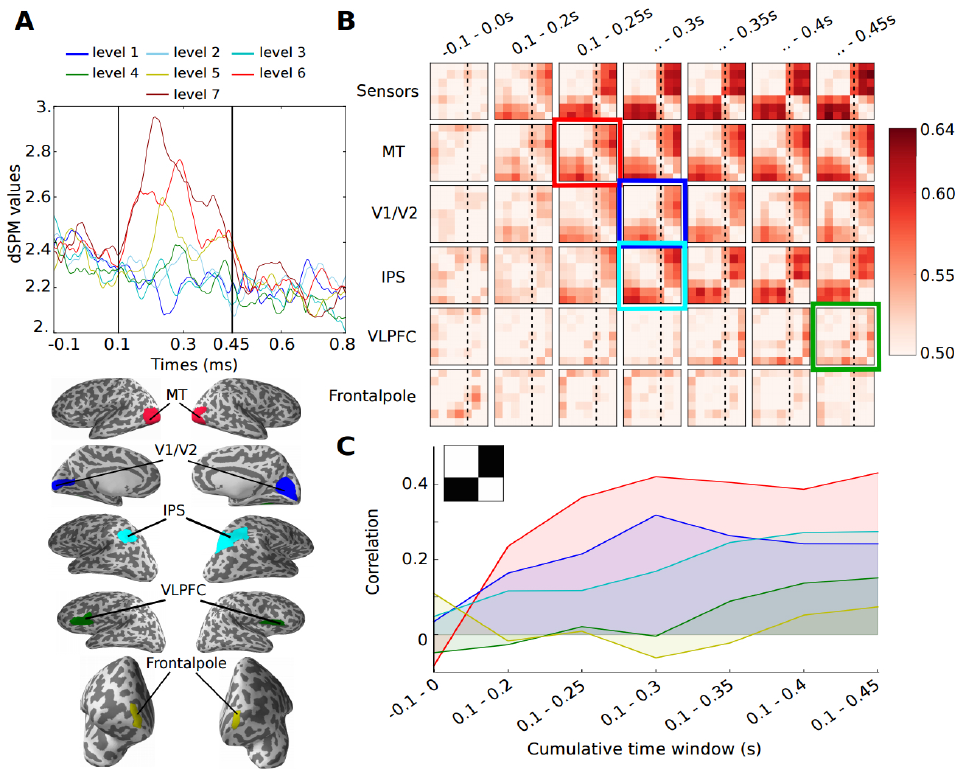
Temporal-accumulation decoding. A) Grand average hMT+ time courses in response to the seven motion coherence levels. B) Grand average similarity matrices (n = 36) in sensors, MT, V1/V2, IPS, VLPFC and frontal pole (top to bottom rows, respectively). Incremental decoding of the similarity matrices within the selected time window could be seen. Colored frames indicate the earliest decoding pattern capturing the perceptual thresholds (dashed lines) e.g. 250 ms for MT. C) Each similarity matrix was correlated with the template matrix optimally capturing perceptual thresholds. Correlations were cumulatively performed over the full time course of brain responses

Using reverse-inference, we selected the template matrix which corresponded to the known mean perceptual thresholds of the 36 participants. We then computed the correlations in specific cortical regions to capture an anatomic and temporal discrimination. The correlation scores between the perceptual templates and the similarity matrices in the different cortical regions are provided in Fig. 2-C. The stability of the similarity matrices (Fig. 2-B) and the plateau of correlations between the template and the similarity matrices (Fig. 2-C) were first reached in hMT+ followed by V1/V2 in occipital regions, IPS and VLPFC. The latency of optimal decoding was consistent with seminal neurophysiology work suggesting functional selectivity of motion computation in hMT+ which may also be indicative of behavioral choice boundary^3, 4, 16, 40^. Perceptual boundaries for motion coherence discrimination could also be decoded later on in regions implicated in the task (V1/V2, IPS and much later in VlPFC) but not in the control region. These observations suggest that the decoder was anatomically and temporally selective. Specifically, the sequence of decoding latencies suggests that the outcome of categorization computed in hMT+ may be forwarded downstream to V1/V2 – as a possible general mechanism contributing to plasticity - as well as VLPFC, as a likely consequence of perceptual decisions required by the task. Decision-related aspect was likely not encoded in low-level sensory areas, however the categorization pattern was still visible in hMT+ when appearing in VLPFC due to accumulation of evidence over the whole time range.

## Discussion

In this study, we showed that brain decoding could classify brain states as a function of visual motion coherence during a discrimination task. The categorical boundary partitioning two brain states was consistent with participants’ discrimination performance as indexed by their perceptual thresholds. Specifically, while the decoder was at chance level in discriminating between two motion coherence levels within the same perceptual category (within perceived or within non-perceived levels of visual motion coherence), the decoder performed well in discriminating brain activity in response to motion coherence levels across different categories (across perceived and non-perceived levels of visual motion coherence). We discuss below the implications and limitations of our findings.

In the visual motion coherence discrimination task used here, the intermixed clouds of dots (or RDKs) were identifiable by two distinct parameters: their color (red or green) and the increased degree of motion coherence in one cloud as comparedto the other. The task required participants to identify the colour of the most coherent cloud of dots. Although the employed stimuli were quite typical for visual motion tasks, a couple requirements set this task apart. First, the selective feature in the display was the coherence of motion irrespective of the direction of motion. This differed from feature-based attention tasks in which the relevant feature is the direction of coherent motion^41, 42^. Second, the task required the discrimination of two clouds of dots simultaneously presented and spatially intermingled; this was distinct from a previous decoding study in which the two populations were spatially segregated^16^. Nevertheless, and consistent with prior decoding work on visual motion processing^5, 15–18^ the earliest robust decoding of motion coherence was found in hMT+, as well as V1/V2. Third, the color of the most coherent cloud of dots was randomized on every trial; as such, the color feature was orthogonal to the task requirement although participants’ effectively classified their responses as”red” or”green”. Accordingly, the successful decoding for any given pair of RDK coherence levels reported here (cf. cells in the decoding matrix being > 50%) captured information about motion coherence *per se*, not its color nor its direction.

The behavioral discrimination of continuous sensory information, such as coherent motion, requires the setting up of an internal criterion classifying sensory information into two or more categories^3, 40^. Seminal work has shown that visual motion coherence at which neural activity reaches 50% of its maximum value can be estimated by means of a neurometric threshold^2^. A similar approach has been used on MEG source estimates in this task, revealing the extent to which the neurometric thresholds computed in the local brain area hMT+ could effectively reflect participants’ discrimination of visual motion coherence^26^. While perceptual thresholds can be derived using several analytical steps and fitting procedures, we have shown that a multivariate decoder can directly capture the partitioning of brain activity as a function of participants’ performance by using a dedicated ranking metric associated with a template matrix correlated with the errors of the decoder when evaluated on left out test data. Our approach also showed that the partitioning of brain states fitting participants’ perceptual thresholds could be found at different timings and at different cortical locations.

Additionally, we found that the more sensory evidence accumulated over time, the more stable and robust the similarity matrices became both at the scalp level and in brain regions. The first stable decoding pattern emerged in hMT+ (~250 ms), consistent with the known likelihood estimations and evidence accumulation of visual motion in this region and at this latency^2, 3^. By 300 ms, a comparable decoding pattern was found in V1/V2, followed by IPS and by 450 ms by VLPFC. The early decoding latencies found in posterior regions and the later latencies found in frontal regions were overall consistent with decoding accuracies reported in perceptual decoding studies. Visual awareness can typically be decoded early in occipital regions and late in frontal areas^20, 43–46^ and while the late decoding component is related to perceptual awareness, it can alsor reflect expectation, task requirements, and attentional selection^20, 43^.

The observed spatiotemporal sequencing and stabilization of peak decoding in regions implicated in the task (but not others, i.e. control area frontal pole) suggest that motion selectivity and choice probability computed in hMT+ could be passed on downstream to early visual cortices as well as to decision-related areas (IPS). Recent models of visual motion processing^4^ and recent fMRI data^22^ have suggested that perceptual priors in early visual cortices may be shaped on the basis of higher-levels computations. Both seminal and recent findings have suggested that attention and feature-selectivity may be crucial in the modulation of early sensory cortices^5, 16, 42^. Our MEG decoding results add to this literature by suggesting that selectivity to higher-order features computed in hMT+ such as motion coherence irrespective of direction or color may feedback to early visual cortices. These and other^17, 18^ results also suggest that the classification boundaries computed in hMT+ may have lasting effects for the analysis of visual motion. In particular, and consistent with previous literature^5, 15–18^, the latency of the categorization pattern across brain regions suggest the possibility that information relevant to perceptual boundaries from hMT+ feedbacks to V1/V2 consistent with predictive coding models of visual processing^4, 47^ and learning theories^48–51^.

Nevertheless, it is noteworthy that in the context of perceptual categorization tasks such as the one employed here, the dissociation between the perceptual and the decisional components are difficult to disentangle^20, 52, 53^. Several studies have discussed the dissociation between perceptual processing and decision-making^20, 21, 52, 54–59^. For instance, a temporal dissociation between early sensory processing in occipital areas and decision-related processing in parieto-frontal regions have been shown to be increasingly pronounced over time^20^. The perceptual thresholds used here to model the best fitting category do not readily dissociate between these two possibilities. Although the present study suggests that multivariate decoding can successfully retrieve perceptual thresholds, it is important to remain skeptical about the link between the information allowing decoding neural activity and its relationship to the computations effectively used to perform the task. For instance, brain activity categorized early on in hMT+ may contain top-down information feedback from decisional brain regions that may have helped the decoded categorization boundaries. However, three main aspects suggest that the decisional component may not be implicated here: first, the decision was made on the orthogonal feature color which was not used in the classifier as reported above. Second, the decoding in parietal cortices occurred much later than the stabilization observed in hMT+. Although response-locked analyses^52^ could be used to disentangle the perceptual and decisional component, one limitation of the current decoder is that it is sensitive to any statistical differences in amplitude or in latency. Hence, analyzing the same time window sorted on the basis of the stimulus onset or of the response would not allow to draw stronger conclusions regarding the (perceptual or decisional) nature of the cortical representations enabling the categorization of brain states. Third, recent evidence suggests that the inactivation of parietal regions are not decisive for motion categorization in monkeys^60^.

In sum, we presented a new MEG decoding technique that can capture the perceived categorization of continuous sensory information. Our results showed a sustainable pattern over time that correlated with participants’ perceptional threshold and which successively implicated hMT+, V1/V2, IPS and VLPFC, consistent with general models of decision-making in motion categorization tasks^61^. Future work should aim at disentangling the perceptual analysis and the decisional components of perceptual decision-making tasks.

## Acknowledgements

This work was supported by an ERC-YStG-263584 and an ANR10JCJC-1904 to V.vW. The funders had no role in study design, data collection and analysis, decision to publish, or preparation of the manuscript. Preliminary results of this work was presented by V.vW. at the Synaesthesia in Perspective workshop (Hamburg, Germany, 2014), by Y. B. at Pattern Recognition in NeuroImaging (Tubingen,¨ Germany, 2014) and by A. G. at Biomag conference (Halifax, Canada, 2014).

## Author contributions statement

N. Z., and V.vW. designed research; Y. B, A. G, N. Z., and V.vW. performed research; Y. B., A. G., and V.vW. contributed unpublished reagents/analytic tools; Y. B., A. G., and V.vW. analyzed data; Y. B., A. G., and V.vW. wrote the paper.

## Additional information

No competing financial interests.

